# A chromosome-level, haplotype-resolved genome assembly of the impressed tortoise, *Manouria impressa*

**DOI:** 10.1101/2025.03.10.642296

**Authors:** Jian Wang, Qinghui Li

**Author notes:** Correspondence: Qinghui Li.

## Abstract

**Background:** The impressed tortoise, *Manouria impressa* (Günther, 1882), is an Endangered, largely fungivorous tortoise of the montane forests of Southeast Asia and one of the two living species of the basal testudinid genus *Manouria*. It is one of the least-studied tortoises, and no chromosome-level genome was available for the genus.

**Findings:** From 68.5 Gb of PacBio HiFi reads (28×), 9.8 Gb of Hi-C read pairs and RNA-seq of heart and lung from a single female, we assembled a 2.42 Gb primary assembly with a contig N50 of 86.9 Mb and a scaffold N50 of 138.6 Mb. Hi-C scaffolding followed by quantitative curation — every scaffold join scored against the within-contig contact-decay expectation — placed 94.5 % of the sequence in 26 chromosomes, matching the testudinid karyotype (2n = 52); the remaining 335 scaffolds are unplaced. Consensus accuracy is QV 67.9 (Merqury), k-mer completeness 95.1 %, and 99.3 % of vertebrate BUSCO genes are complete. Two phased haplotype assemblies (scaffold N50 87.8 and 91.6 Mb; QV 69.1 and 68.5; 99.2 % combined k-mer completeness) and the 18.1-kb mitochondrial genome were assembled from the same data. Repeats mask 48.6 % of the genome (LINEs 11.5 %, LTR retrotransposons 10.0 %, DNA transposons 7.6 %, SINEs 2.6 %, unclassified families 15.4 %); 24,875 protein-coding genes were predicted from RNA-seq and OrthoDB protein evidence (97.6 % complete vertebrate BUSCOs), 89.0 % of them functionally annotated.

**Conclusions:** This haplotype-resolved reference, with its curation evidence and scripts, provides the first chromosome-level genomic resource for *Manouria* and a foundation for conservation genomics and for studies of the dietary specialisation of this fungivorous tortoise.

## Context

The impressed tortoise (*Manouria impressa*) is a medium-sized tortoise of montane evergreen forest from southern China and Myanmar through Thailand, Laos, Cambodia and Vietnam, recognisable by its flattened, serrated, golden-brown carapace with concave ("impressed") scutes [65,66,71]. It is unusual among tortoises in its diet, which consists largely of fungi — it is regarded as a near-obligate mycophage [72,73] — supplemented by bamboo shoots and other plant material [65,69,70]; through this diet it is thought to contribute to nutrient cycling and fungal spore dispersal in its forest habitat [72]. It is secretive and seasonally inactive [69,70]. The species is listed as Endangered on the IUCN Red List [66] and in CITES Appendix II (family listing Testudinidae spp.) [75]; populations are declining through habitat loss and collection for the pet and food trades, a low reproductive rate and narrow habitat requirements make recovery slow, and the species is difficult to keep and breed in captivity [66,67,71,73]. It remains one of the least-studied tortoises, with little ecological and no genomic data to support conservation planning [66]. *Manouria* (two species: *M. impressa* and the Asian forest tortoise *M. emys*) together with the North American *Gopherus* forms the most basal lineage of living Testudinidae [76], so its genome also anchors the base of the tortoise radiation.

Reference genomes now exist for several turtle lineages — the painted turtle *Chrysemys picta* [48], the soft-shelled turtle *Pelodiscus sinensis* and green sea turtle *Chelonia mydas* [49], the red-eared slider *Trachemys scripta* [53] and the pond turtles *Mauremys reevesii* and *M. mutica* [59,60] — and, within Testudinidae, for Agassiz’s desert tortoise *Gopherus agassizii* [50,51] and Morafka’s desert tortoise *G. morafkai* [57], Goode’s thornscrub tortoise *G. evgoodei* (Vertebrate Genomes Project assembly rGopEvg1 [23,52]), the Galápagos giant tortoise *Chelonoidis abingdonii* [54,56], the Aldabra giant tortoise *Aldabrachelys gigantea* [55] and the spur-thighed tortoise *Testudo graeca* [58]. Of these, only *G. evgoodei*, *A. gigantea* and *G. morafkai* are chromosome-level assemblies built from long reads with Hi-C; *A. gigantea* [55] is the closest methodological precedent (PacBio HiFi + Hi-C). Testudinidae share a conserved karyotype of 2n = 52 [61–63]. No genome of *Manouria* has been described in the literature.

Here we present a chromosome-level, haplotype-resolved reference genome of *M. impressa* assembled from PacBio HiFi and Hi-C data of a single female, with a quantitative Hi-C curation record, two phased haplotype assemblies, the mitochondrial genome and an RNA-seq-supported gene annotation, built with a fully scripted and logged pipeline so that every number reported here can be regenerated from the raw reads.

## Methods

### Sample

All data derive from a single adult female *M. impressa* (individual ID **IT-1**; BioSample SAMC4973488 for the liver DNA libraries, SAMC4973532 and SAMC4976828 for the heart and lung RNA libraries), estimated age about 20 years, collected in Hekou, Yunnan, China, in 2013 under permit from the Yunnan Provincial Forestry Department (Yunlin Conservation Permit [2013] No. 86) and the Yunnan Daweishan National Nature Reserve, and kept in captivity until it died of natural causes on 29 June 2023. The carcass was placed in a −80 °C freezer immediately on discovery and thawed the following day for dissection; liver, heart and lung were aliquoted into individual tubes, returned to −80 °C for one month, and shipped to the sequencing provider (Annoroad Gene Technology, Beijing, China) on dry ice (3 days). Liver was used for high-molecular-weight DNA (HiFi sequencing) and for the Hi-C library; total RNA was extracted from heart (library IT3) and lung (library IT4). No procedure was performed on a living animal; the study was approved by the Academic Ethics Committee of the School of Life Sciences, Hainan Normal University (Approval No. 202532).

### Sequencing

High-molecular-weight DNA was sequenced on a PacBio **Revio** system in four SMRT cells (movies m84069_230831, m84129_231016, m84129_231026, m84129_231103; circular consensus by ccs 7.0.0), yielding 3,618,287 HiFi reads (68.55 Gb; mean 18,945 bp, N50 18,784 bp, maximum 65,258 bp), all with predicted read accuracy ≥ Q20. The Hi-C library was prepared from liver with the restriction enzyme MboI and sequenced on an MGI DNBSEQ platform (2 × 150 bp), yielding 32,770,909 read pairs (9.83 Gb); 55 % of reads carry the MboI ligation junction GATCGATC. Poly(A)-selected RNA-seq libraries from heart and lung were sequenced as 2 × 150 bp reads on an MGI DNBSEQ-G400 (platform inferred from the read-name format): 22,319,011 and 22,448,804 read pairs (6.70 and 6.73 Gb). All read counts were recomputed from the raw files (Table 2). No short-read whole-genome data were used.

**Table 1.** Software, versions and non-default parameters.

| Tool | Version | Used for | Non-default parameters |
|---|---|---|---|
| samtools | 1.21 | read preparation, Hi-C mapping/scaffolding |  |
| seqkit | 2.9.0 | read preparation, release files |  |
| meryl | 1.4.1 | read preparation, assembly QC | k = 21 |
| genomescope2 | 2.0 | read preparation | -k 21 -p 2 |
| hifiasm | 0.25.0 | contig assembly | --h1/--h2 (Hi-C mode) |
| gfatools | 0.5.5 | contig assembly |  |
| chromap | 0.2.6 | Hi-C mapping/scaffolding | --preset hic -q 1 --remove-pcr-duplicates |
| yahs | 1.2.2 | Hi-C mapping/scaffolding | -e GATC |
| minimap2 | 2.28 | synteny, coverage, release files | -x asm20 (synteny), map-hifi (coverage), asm5 (haplotypes, v2.4) |
| pretextmap | 0.1.9 | contact maps |  |
| pretextsnapshot | 0.0.4 | contact maps |  |
| bedtools | 2.31.1 | Hi-C statistics |  |
| busco | 5.8.0 | assembly QC, functional / ncRNA annotation | vertebrata_odb10, offline |
| merqury | 1.3 | assembly QC | k = 21 |
| assembly-stats | 1.0.1 | assembly QC |  |
| diamond | 2.2.8 | functional / ncRNA annotation |  |
| eggnog-mapper | 2.1.12 | functional / ncRNA annotation | -m diamond --tax_scope 7742 --go_evidence non-electronic |
| compleasm | 0.2.10 (bioconda package 0.2.6) | assembly QC | vertebrata_odb12, sauropsida_odb12 |
| HapHiC | commit 74f1cb2 | haplotype scaffolding | pipeline --RE GATC, 26 groups |
| juicer_tools | 1.22.01 | Hi-C mapping/scaffolding | pre |
| BRAKER3 | 3.0.8 | gene prediction | --bam (HISAT2) --prot_seq OrthoDB v12 Vertebrata --busco_lineage vertebrata_odb10 |
| RepeatModeler | 2.0.5 | repeat library | -LTRStruct |
| RepeatMasker | 4.1.7-p1 | repeat masking | -xsmall; species library then -species tetrapoda |
| FCS-adaptor | 0.5.4 | contamination / mitogenome | --euk |
| FCS-GX | v0.5.5 | contamination / mitogenome | --tax-id 74924 |
| MitoHiFi | MitoHiFi 3.2.1 | contamination / mitogenome | -f/-g NC_011815.1 -o 2 |
| tRNAscan-SE | 2.0.12 | tRNA annotation | -E -H --detail; EukHighConfidenceFilter |
| Barrnap | 0.9 | rRNA annotation | --kingdom euk |
| paftools.js | minimap2 2.28 | haplotype divergence | call -l 10000 -L 50000 |
| TSEBRA | BRAKER 3.0.8 | gene-set merge (compared, not used for the final set) | default |
| bedtools | 2.31.1 | RNA-seq support per gene; soft-mask union | multicov; maskfasta -soft |

**Table 2a.**
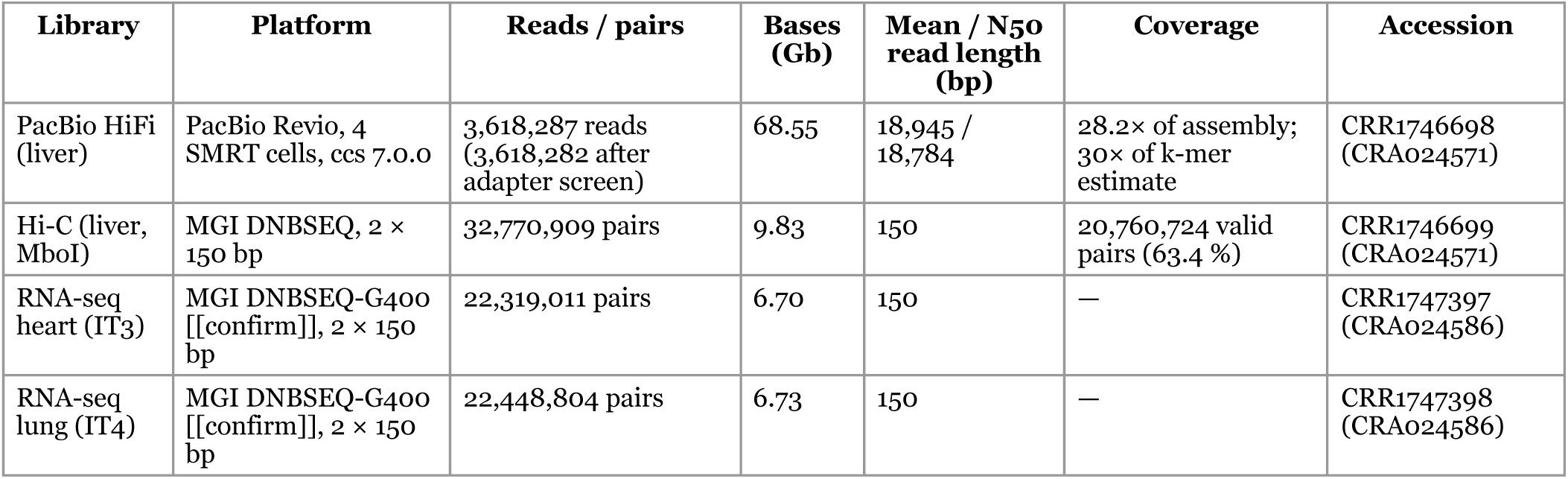
Sequencing data (counted from the raw files)

| Library | Platform | Reads / pairs | Bases (Gb) | Mean / N50 read length (bp) | Coverage | Accession |
|---|---|---|---|---|---|---|
| PacBio HiFi (liver) | PacBio Revio, 4 SMRT cells, ccs 7.0.0 | 3,618,287 reads (3,618,282 after adapter screen) | 68.55 | 18,945 / 18,784 | 28.2× of assembly; 30× of k-mer estimate | CRR1746698 (CRA024571) |
| Hi-C (liver, MboI) | MGI DNBSEQ, 2 × 150 bp | 32,770,909 pairs | 9.83 | 150 | 20,760,724 valid pairs (63.4 %) | CRR1746699 (CRA024571) |
| RNA-seq heart (IT3) | MGI DNBSEQ-G400 [[confirm]], 2 × 150 bp | 22,319,011 pairs | 6.70 | 150 | — | CRR1747397 (CRA024586) |
| RNA-seq lung (IT4) | MGI DNBSEQ-G400 [[confirm]], 2 × 150 bp | 22,448,804 pairs | 6.73 | 150 | — | CRR1747398 (CRA024586) |

**Table 2b.** Assembly statistics.

| **Statistic** | **Mimp_v3 (reference)** | **hap1** | **hap2** |
|---|---|---|---|
| Sequences | 361 | 414 | 396 |
| Total length (bp) | 2,417,402,780 | 2,365,101,262 | 2,380,510,770 |
| Scaffold N50 (bp) | 138,646,552 | 87,800,537 | 91,606,729 |
| Scaffold N90 (bp) | 21,674,238 | 17,392,175 | 18,583,814 |
| Longest sequence (bp) | 382,984,538 | 146,557,687 | 141,836,076 |
| Contigs | 509 | 638 | 624 |
| Contig N50 (bp) | 86,871,185 | 36,595,271 | 28,880,847 |
| Gaps | 148 | 224 | 228 |
| Chromosome-scale sequences | 26 (94.5 % of length) | — | — |
| Merqury QV / k-mer completeness (%) | 67.93 / 95.09 | 69.12 / 94.47 | 68.51 / 94.78 (both haplotypes: QV 68.80, completeness 99.18) |
| compleasm vertebrata_odb12 (n = 3,390) (%) | S 98.82 D 0.94 F 0.12 M 0.12 | S 98.67 D 0.68 F 0.15 M 0.50 | S 99.00 D 0.74 F 0.12 M 0.15 |
| BUSCO 5.8.0 vertebrata_odb10 (n = 3,354) (%) | C 99.3 [S 97.8, D 1.5] F 0.3 M 0.4 | C 98.7 [S 97.4, D 1.3] F 0.4 M 0.9 | C 99.0 [S 97.8, D 1.2] F 0.4 M 0.6 |
| EBP quality notation | 7.8.Q67 | 7.7.Q69 | 7.7.Q68 |
| Accession | GWHHIVU00000000.2 [[PENDING]] | [[PENDING]] | [[PENDING]] |

**Table 3.** Annotation statistics (final evidence-supported gene set)

| **Feature** | **Value** |
|---|---|
| Protein-coding genes | 24,875 |
| Transcripts | 29,927 |
| Transcripts per gene | 1.20 |
| Mean gene length (bp) | 33,694 |
| Mean CDS length (bp) | 1,668 |
| Mean exons per transcript | 9.82 |
| Mean exon / intron length (bp) | 170 / 4,418 |
| Median protein length (aa) | 380 |
| Single-exon transcripts | 4,355 |
| BUSCO vertebrata_odb10, protein mode (n = 3,354) (%) | C 97.6 [S 95.8, D 1.9] F 1.4 M 1.0 |
| Genes with eggNOG-mapper annotation (%) | 22,136 (89.0); GO 15,241, KEGG 15,259, Pfam 20,957 |
| Genes with RNA-seq support (heart + lung) | 18,875 (75.9 %) with ≥ 10 reads; 22,490 (90.4 %) with ≥ 1 read |
| tRNA genes (tRNAscan-SE high-confidence / all predictions) | 3,309 / 24,407 |
| rRNA genes or fragments (Barrnap) | 18S 92; 5.8S 79; 28S 99; 5S 181 |
| Repeats (soft-masked, union of two passes) (%) | 48.60 |

### Read preparation and genome survey

HiFi reads were extracted from the BAM files with samtools 1.21 [14], filtered to predicted accuracy rq ≥ 0.99 (all reads passed), and screened for residual SMRTbell adapter sequence with SeqKit 2.9.0 [15,16]; five reads containing adapter were removed (3,618,282 reads retained). 21-mers were counted with meryl 1.4.1 and the k-mer spectrum modelled with GenomeScope 2.0 [11] (ploidy 2), giving a haploid genome size of 2.26 Gb and heterozygosity 0.31–0.35 %; the model assigns 597 Mb (26.4 %) of the haploid genome to k-mers present in more than one copy ("Genome Repeat Length"), a k-mer-level quantity that is not comparable with homology-based repeat annotation (see Data validation). HiFi coverage is 28.2× of the final assembly (30× of the k-mer estimate).

### Contig assembly

Contigs were assembled with hifiasm 0.25.0 [1,2] in Hi-C-integrated mode (`hifiasm --h1 R1 --h2 R2`), default parameters, which produces a primary assembly and two Hi-C-phased haplotype assemblies (hap1, hap2). The primary assembly comprised 460 contigs (2,428,913,094 bp, N50 86,871,185 bp); hap1 and hap2 comprised 686 and 631 contigs (N50 36.6 and 28.9 Mb).

### Hi-C scaffolding

Hi-C [43,44] reads were aligned to each assembly with chromap 0.2.6 [5] (`--preset hic -q 1 -- remove-pcr-duplicates`). Of 32,770,909 pairs, 84.5 % of reads mapped, 70.6 % uniquely; after duplicate removal and MAPQ ≥ 1 and proper-pair filtering, 20,760,724 valid pairs (63.4 %) were used for scaffolding, 48.4 % of them intra-contig (26.3 % at ≥ 20 kb). The primary assembly was scaffolded with YaHS 1.2.2 [3] (`-e GATC`), yielding 394 scaffolds (165 joins; scaffold N50 138.6 Mb; 26 chromosome- scale scaffolds). The haplotype assemblies were scaffolded with HapHiC (commit 74f1cb2) [4] (`haphic pipeline --RE GATC`, 26 groups). HapHiC was also run on the primary assembly (scaffold N50 106.2 Mb) and the YaHS result was retained for the primary because it recovered more complete chromosomes; both are reported. Contact maps were generated with PretextMap 0.1.9 [8] and Juicer Tools 1.22.01 [6], and inspected in Juicebox Assembly Tools 1.11.08 [7].

### Quantitative curation

Instead of relying on visual editing alone, every scaffold join was scored from the Hi-C alignments (script `join_qc.py`): contacts across each junction (100-kb bins, 2-Mb windows) were expressed (i) as a ratio to the expectation for contiguous sequence, derived from the within-contig contact-distance decay, and (ii) as a ratio to the inter-scaffold background. The scaffolder’s own 165 joins calibrated the statistic (background ratio median 46, 5th percentile 11); a junction independently verified by eye in Juicebox scored 0.62 / 45. Joins were broken when the background ratio was < 5 and candidate adjacencies between scaffolds were accepted when ≥ 11 (`curation_contacts.py`, `apply_curation.py`). Whole- genome alignment to the *G. evgoodei* reference rGopEvg1 (GCF_007399415.2) with minimap2 2.28 [12,13] (`-x asm20`, alignments ≥ 50 kb, MAPQ ≥ 30) was used only to prioritise what to examine; Hi-C decided. Three breaks were made (a circular 6.08-Mb satellite contig spliced into a chromosome; two unsupported joins in small scaffolds) and three joins (1.5-, 1.1- and 1.9-Mb scaffolds attached to chromosome ends with background ratios 59, 35 and 15); four synteny-suggested joins were declined (ratios 7.7–8.6). One apparent fusion relative to *G. evgoodei* (v3 chr12 = GopEvg chr14 + chr21) showed continuous contact across the seam (6.6× background) and was kept as a genuine difference. After editing, Hi-C reads were re-aligned and all junctions re-scored (new junctions 58 / 39 / 16; no junction < 5). Every edit is listed with its evidence in Supplementary Table S2.

### Chromosome naming and release

The 26 chromosome-scale scaffolds were named chr1–chr26 in order of decreasing length and oriented to match *G. evgoodei* by a length-weighted strand vote of the alignments; unplaced scaffolds were named scaffold_1…scaffold_368 by length (`name_chromosomes.py`). After removal of contaminant scaffolds (below), the release (Mimp_v3) comprises 361 sequences, 2,417,402,780 bp, 509 contigs (contig N50 86,871,185 bp), scaffold N50 138,646,552 bp, longest chromosome 382,984,538 bp, 148 gaps of 100 N (proximity-ligation gaps of unknown size; AGP provided). Relative to *G. evgoodei* (24 named chromosomes), chr1–chr11 are one-to-one with GopEvg chr1–chr11; GopEvg chr3 corresponds to v3 chr3 + chr22; two microchromosomes (chr17, chr25) are repeat-rich and largely unalignable (Figure 1B; alignment in Figure 2D).

**Figure 1.**
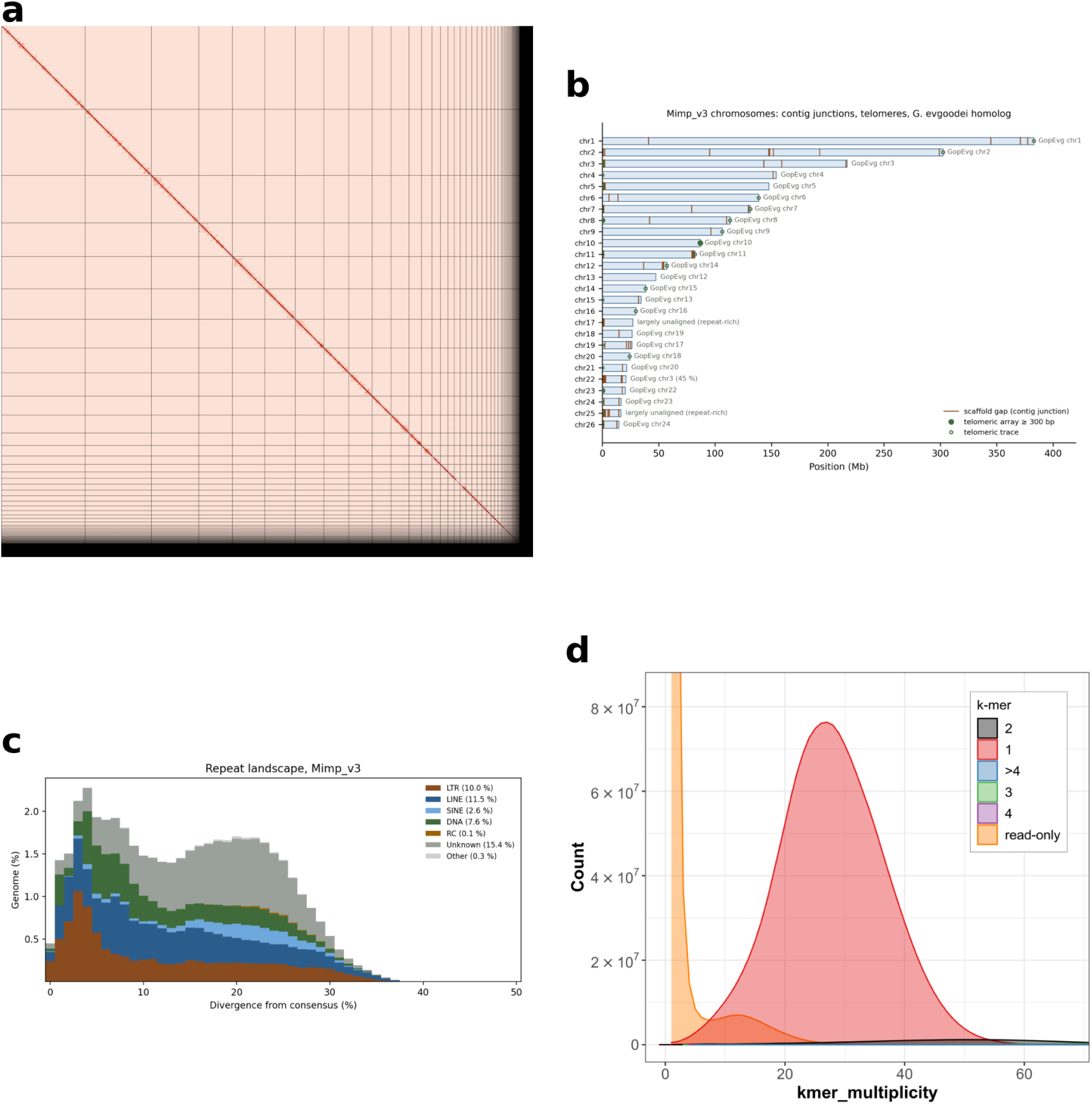
Overview of the Mimp_v3 assembly. (A) Hi-C contact map of the release (chr1–chr26, then unplaced scaffolds). (B) The 26 chromosomes with contig junctions, telomeric arrays and the *Gopherus evgoodei* homolog of each. (C) Repeat landscape (genome fraction per 1 % divergence bin, by class). (D) Merqury copy-number spectrum (QV 67.9, k-mer completeness 95.1 %).

**Figure 2.**
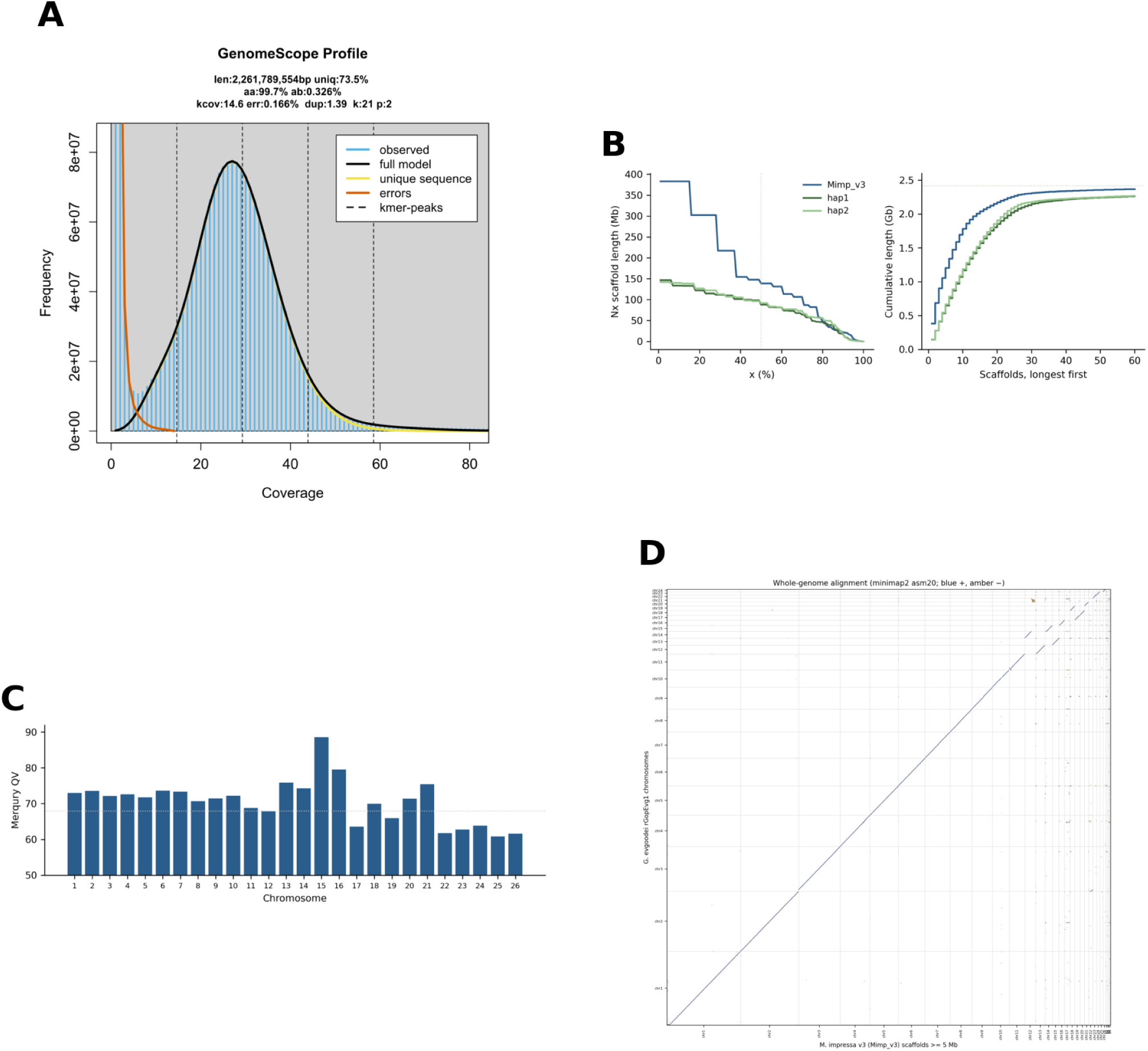
Assembly validation. (A) GenomeScope 2.0 21-mer spectrum of the HiFi reads (haploid length 2.26 Gb, heterozygosity 0.33 %). (B) Nx curves and cumulative length for Mimp_v3, hap1 and hap2. (C) Merqury QV per chromosome (dotted line: genome-wide QV). (D) Whole-genome alignment of Mimp_v3 to *G. evgoodei* rGopEvg1 (minimap2 asm20, alignments ≥ 50 kb, MAPQ ≥ 30).

The two haplotype assemblies were released as scaffold-level alternate assemblies: scaffolds renamed by length (h1_scaffold_N, h2_scaffold_N), sequences < 1 kb removed, and each scaffold assigned to its best-matching primary chromosome by minimap2 (`asm5`) (Supplementary Table S6). They are not chromosome-named because, at 0.3 % heterozygosity, phasing is in blocks and the 26 HapHiC groups are not guaranteed to be whole chromosomes.

### Assembly quality control

Consensus quality and k-mer completeness were estimated with Merqury 1.3 [10] against the HiFi 21- mer database; gene completeness with compleasm 0.2.10 [19] (vertebrata_odb12, n = 3,390; sauropsida_odb12, n = 6,038) and BUSCO 5.8.0 [17,18] (vertebrata_odb10, n = 3,354). HiFi reads were re-aligned to the release (minimap2 `map-hifi`) and depth computed in 10-kb windows; zero-coverage runs on chromosomes were examined by counting reads at all mapping qualities and aligning the region to the alternate haplotype. Telomeric repeats ((TTAGGG)n/(CCCTAA)n) were counted in the terminal 50 kb of every chromosome. Haplotype divergence was measured by aligning hap1 to hap2 and each to the primary (minimap2 `asm5 --cs`; `paftools.js call`, blocks ≥ 10 kb) . Contamination was screened with NCBI FCS-adaptor 0.5.4 and FCS-GX 0.5.5 [21] (taxid 74924) on all three nuclear assemblies.

### Mitochondrial genome

The mitogenome was assembled from the HiFi reads with MitoHiFi 3.2.1 [22] using the published *M. impressa* mitogenome NC_011815.1 (GenBank EF661586.1, direct submission [74]) as reference (`-o 2`), and the three nuclear assemblies were screened for mitochondrial sequence by alignment of the mitogenome (minimap2 `asm10`, MAPQ ≥ 30; sequences ≥ 80 % covered would be removed — none were).

### Repeat annotation

A species-specific repeat library was built with RepeatModeler 2.0.5 [26] (`-LTRStruct`) in the Dfam TETools container (2,492 families: 414 LTR, 288 LINE, 276 DNA, 94 SINE, 1,375 unclassified). The release was soft-masked with RepeatMasker 4.1.7-p1 [27] in two passes — the species library, then the Dfam 3.8 [28] Tetrapoda library on the masked output — and the two annotations were merged. The species-library pass masked 48.34 % of the assembly; the Dfam pass annotated a further 0.84 %, partly overlapping the first. Because RepeatMasker uppercases its input, the final soft-masked genome was built as the union of the two annotations (bedtools maskfasta), and all repeat fractions reported here are that union over the cleaned assembly: 1,174,831,695 of 2,417,402,780 bp = **48.60 %** (the per-class values are from the species-library pass summary, which covers 48.34 %). A repeat landscape (divergence from consensus per class, 1 % bins) was computed from the RepeatMasker output with class totals taken from the `.tbl` summary.

### Gene annotation

RNA-seq reads from heart and lung were aligned to the soft-masked genome with HISAT2 2.2.1 [34] (`-- dta`; 97.1 % and 97.7 % overall alignment). Protein-coding genes were predicted with BRAKER 3.0.8 [29] using the RNA-seq alignments and the OrthoDB v12 Vertebrata protein set [20] as evidence (GeneMark-ETP [30] and AUGUSTUS 3.5 [31,32], combined with TSEBRA [33]). BRAKER3 yields three gene sets — GeneMark-ETP, AUGUSTUS predicted with RNA-seq and protein hints, and their TSEBRA merge — and each was assessed with BUSCO in protein mode on the longest isoform per gene: TSEBRA 17,392 genes (C 89.6 %), GeneMark-ETP 27,692 (C 90.2 %, D 8.5 %), AUGUSTUS 26,279 (C 97.6 %, D 1.9 %); a union of the TSEBRA set with non-overlapping AUGUSTUS genes scored 96.2 %, and a TSEBRA re-run keeping all AUGUSTUS transcripts 97.8 % with 19.1 % duplication. By a rule fixed before the comparison (highest completeness with duplication below 4 %), the AUGUSTUS set was taken forward and filtered to genes supported by at least one line of evidence — ≥ 1 RNA-seq read over the gene span, an eggNOG-mapper ortholog hit, or a BUSCO match — which removed 1,404 models. Genes were renamed MIMP3_gNNNNN in chromosome order, and the GFF3 was checked for features outside sequence bounds and duplicate identifiers (none). The longest isoform per gene was functionally annotated with eggNOG-mapper 2.1.12 [35] (DIAMOND [37] search of eggNOG 5.0 [36], taxonomic scope Vertebrata, GO terms from non-electronic evidence; Pfam [42], KEGG [41] and GO [40] assignments). tRNA genes were predicted with tRNAscan-SE 2.0.12 [38] (eukaryotic model) and filtered with its EukHighConfidenceFilter to remove tRNA-derived SINE copies and pseudogenes; rRNA genes were predicted with Barrnap 0.9 [39]. Statistics of the final set are in Table 3.

### Chromosome correspondence and structural differences relative to *G. evgoodei*

Alignments of Mimp_v3 to rGopEvg1 (minimap2 `asm20`, ≥ 50 kb, MAPQ ≥ 30) were chained into synteny blocks (same reference chromosome and strand, gaps < 3 Mb) and blocks ≥ 1 Mb were retained. Candidate structural differences were defined as adjacent blocks on a Mimp_v3 chromosome that map to different reference chromosomes, switch strand, or jump ≥ 5 Mb in reference order.

Because such differences can equally reflect an error in either assembly, each candidate breakpoint was tested on the Mimp_v3 side: whether it lies inside a single HiFi contig, and the Hi-C contact across it (2-Mb windows on each side) relative to the within-contig distance-decay expectation and to the inter- chromosomal background, using the same statistic as the curation (16_synteny_rearrangements.py).

Differences are reported only where Mimp_v3 is physically continuous (or, for separate chromosomes, where Hi-C contact between them is at background); the direction of change between the two species is not inferred.

### Correspondence to the preliminary version

A preliminary assembly of the same data had been deposited as version .1 of the Genome Warehouse record. For users of that version, every one of its sequences was aligned to Mimp_v3 (minimap2 `asm5`, alignments ≥ 50 kb, MAPQ ≥ 30) and assigned to its best-matching v3 sequence (Supplementary Table S4).

### Software and reproducibility

All steps are Slurm scripts with pinned software versions (Table 1; `software_versions.tsv` generated from the environments), logged with job identifiers in a protocol file that is the source of this Methods section. The complete pipeline, the curation evidence (join scores, edit log), and the scripts that generate every table and figure are available in the project repository and archived at Zenodo (see Availability of source code).

## Data validation and quality control

**Sequence data.** Read counts, bases, read-length statistics and coverage for each library, recomputed from the raw files, are in Table 2. The HiFi k-mer spectrum is unimodal with a single homozygous peak at 27× and a small heterozygous shoulder (Figure 2A), consistent with the 0.31–0.35 % heterozygosity estimate.

### Assembly contiguity and structure

The primary assembly places 2,284,523,576 bp (94.5 %) of the sequence in 26 chromosomes, matching the Testudinidae karyotype of 2n = 52. The Hi-C contact map of the release shows 26 diagonal blocks with no off-diagonal signal (Figure 1A). Alignment to *G. evgoodei* shows one-to-one correspondence for chr1–chr11 and the expected microchromosome differences (Figure 2D; chromosome summary in Figure 1B).

### Chromosome structure relative to *G. evgoodei*

Twenty-two of the 26 Mimp_v3 chromosomes correspond to a single *G. evgoodei* chromosome over ≥ 1-Mb synteny blocks (Supplementary Table S12): chr1–chr11 to GopEvg chr1–chr11 in order, and chr13–chr16, chr18, chr20, chr21, chr23, chr24 and chr26 to one microchromosome each; chr17 and chr25 are repeat-rich and have no block ≥ 1 Mb. Four structural differences between the two assemblies are supported on the Mimp_v3 side: (i) Mimp_v3 chr12 (57.2 Mb) carries GopEvg chr14 (33.3 Mb) and chr21 (10.0 Mb) in one molecule — the junction lies inside a single HiFi contig (ptg000055l) and Hi-C contact across it is 18× the inter- chromosomal background (with the reduced short-range contact typical of a centromeric region, as noted during curation); (ii) GopEvg chr3 is represented by two Mimp_v3 chromosomes, chr3 (216.7 Mb) and chr22 (8.5 Mb aligned), between which Hi-C contact is at background (0.84×), i.e. they are separate molecules in *M. impressa*; (iii) a 1.5-Mb segment homologous to GopEvg chr19 lies within Mimp_v3 chr2 (165.4–166.8 Mb), both boundaries inside contig ptg000008l; (iv) intra-chromosomal differences — a strand switch at 7.4 Mb and an 8-Mb order jump at 10.3 Mb on chr11, a 5.4-Mb order jump on chr12 and a strand switch at 21.9 Mb on chr19 — all lie inside single HiFi contigs with Hi-C contact 28–54× background across the breakpoint. These are therefore features of the *M. impressa* assembly rather than scaffolding errors; whether each represents a rearrangement in the *M. impressa* lineage, in *Gopherus*, or an unresolved join in rGopEvg1 cannot be decided from one genome pair, and the equal chromosome number of the two species (2n = 52) is consistent with one fusion and one fission in either direction. Candidate differences at contig junctions that failed the Hi-C test would have been excluded; none occurred. In the Earth BioGenome Project notation [24,25] the assembly is **7.8.Q67** (contig N50 10⁷ bp, scaffold N50 10⁸ bp, QV 67), above the 6.7.Q40 reference standard.

### Accuracy and completeness

Merqury QV is 67.93 (8,173 erroneous k-mers; estimated error 1.6 × 10⁻⁷) and k-mer completeness 95.09 % (the residual read k-mers absent from the assembly include those of the removed bacterial sequence, which remains in the reads); per-chromosome QV ranges from 60.8 (chr25) to 88.6 (chr15) (Figure 2C). compleasm finds 98.8 % of vertebrata_odb12 genes complete and single-copy (D 0.9 %, F 0.1 %, M 0.1 %) and BUSCO 99.3 % of vertebrata_odb10 genes complete (S 97.8 %, D 1.5 %, F 0.3 %, M 0.4 %) (Table 2; Merqury spectrum in Figure 1D).

### Coverage

Median HiFi depth over the release is 27.8× (243,083 10-kb windows); 222 windows (34 on chromosomes) have < 5 % of median depth and 152 windows > 2.5× median (collapsed repeats or organelle sequence). The two largest zero-coverage runs on chromosomes (chr1:42.87–43.13 Mb and scaffold_28:0.25–0.57 Mb) are heterozygous tandem duplications: reads are present at all mapping qualities (unfiltered depth 18.5×) but map with MAPQ 0 to two near-identical copies, and the region aligns twice to the alternate haplotype. They were left unedited and are flagged for users calling variants there.

### Telomeres

Telomeric arrays ≥ 300 bp are present at 5 of 52 chromosome ends (chr3, chr5, chr8 and chr23 start; chr10 end; longest 1,134 bp), with shorter traces at 20 more and none in unplaced scaffolds (Supplementary Table S5). Short telomeric arrays at scaffold ends are typical of HiFi + Hi-C assemblies of large repeat-rich genomes and indicate that most chromosome ends terminate in subtelomeric repeats rather than in the telomere itself.

### Haplotypes

hap1 (414 scaffolds, 2.365 Gb, N50 87.8 Mb) and hap2 (396 scaffolds, 2.381 Gb, N50 91.6 Mb; Hi-C contact maps in Supplementary Figure S1) have QV 69.1 and 68.5 (68.8 combined); their k- mer completeness is 94.5 % and 94.8 % individually and 99.2 % combined, i.e. together they contain essentially all read k-mers. compleasm single-copy completeness is 98.7 % (hap1) and 99.0 % (hap2) with duplication below 0.8 %, as expected for a haplotype, and BUSCO completeness is 98.7 % and 99.0%. Across 2.21 Gb of aligned sequence the two haplotypes differ by 4,572,296 SNVs (2.07 per kb), 573,150 small indels (0.26 per kb) and 50,175 structural variants ≥ 50 bp — a nucleotide divergence of 0.23 %, consistent with the k-mer heterozygosity estimate; the primary assembly is roughly equidistant from both (1.02 and 0.95 SNVs per kb), as expected for a mosaic. Phasing accuracy could not be measured directly because no parental data exist; the haplotypes should be regarded as block-phased.

### Mitochondrial genome

A single circular contig of 18,133 bp carries the 13 protein-coding genes, 22 tRNAs and 2 rRNAs of the vertebrate mitogenome. It is 1.5 kb longer than NC_011815.1 (16,642 bp) owing to a tandem array of four tRNA-Thr/tRNA-Pro units adjacent to the control region Among 1,529 HiFi reads that span the whole array into the origin, 44 % (677) carry the four-unit structure of the consensus and 52 % (789) carry five units (one ∼300-bp unit more); other copy numbers are rare. The array is therefore length-heteroplasmic; the deposited sequence is the four-unit MitoHiFi consensus.. No mitochondrial sequence remains in the nuclear assemblies.

### Contamination

FCS-adaptor flagged no sequence in any assembly. FCS-GX identified bacterial sequence in 33 unplaced scaffolds of the primary assembly (11.5 Mb; the largest 4.4 Mb), 48 scaffolds plus one 2.9-kb internal segment of hap1 (11.5 Mb) and 7 scaffolds of hap2 (4.6 Mb); no chromosome was affected. The taxa — *Parabacteroides*, *Bacteroides*, *Prevotella*, *Lactococcus*, *Morganella*, *Erysipelothrix* and one phage — are gut and post-mortem flora, as expected for liver sampled from an animal that died naturally, and they account for the high-depth windows noted above. All flagged sequences were removed (the internal segment hard-masked) with `gx clean-genome`, and a second FCS- GX screen of the cleaned assemblies flagged nothing. Supplementary Table S11 lists every removed sequence with its taxonomic assignment. All statistics in this paper refer to the cleaned assemblies.

### Repeats

The union of the two RepeatMasker annotations soft-masks 48.60 % of the cleaned assembly (Table S9); by class (species-library pass, 47.05 % interspersed): LINEs 11.5 % (CR1/L2 9.3 %), LTR retrotransposons 10.0 % (Gypsy 6.2 %, endogenous retroviruses 2.3 %), DNA transposons 7.6 % (hAT 3.4 %, Harbinger 3.0 %), SINEs 2.6 %, rolling-circle elements 0.1 % and 15.4 % unclassified families, with simple repeats, low-complexity sequence and satellites adding 0.6 %. The repeat landscape (Figure 1C) shows a recent burst of LTR elements at 2–5 % divergence from consensus superimposed on an older, broad peak of unclassified and DNA elements at 15–30 %.

The GenomeScope "repeat length" of 597 Mb (26.4 % of the haploid genome) is a different quantity and is not expected to agree with the masked fraction: it is the part of the genome covered by 21-mers that occur more than once in the haploid genome, so repeat copies that have diverged by more than a few percent — the majority of the repeat mass here, which lies at 15–30 % divergence — no longer share identical 21-mers and are counted as unique sequence. Homology-based annotation recognises these diverged copies. The assembly is also 7 % larger than the k-mer genome-size estimate (2.42 vs 2.26 Gb), for the same reason: collapsed repeat copies are under-counted by k-mer models. The two values therefore bracket the repeat content: the k-mer estimate is a lower bound for near-identical repeats, and 48.6 % is the fraction of the assembly assignable to repeat families; we report the latter as the repeat content of the genome.

### Gene annotation

tRNAscan-SE predicted 24,407 tRNA-like sequences, of which 3,309 pass the high-confidence filter (the remainder are pseudogenes and tRNA-derived SINE copies); the high- confidence set covers all 20 amino-acid isotypes plus initiator Met and selenocysteine tRNAs. Barrnap found 92 18S, 79 5.8S, 99 28S and 181 5S rRNA genes or fragments. The final set contains 24,875 protein-coding genes and 29,927 transcripts (1.20 per gene), 23,000 of them on chromosomes (Table 3; Figure 3A). 97.6 % of vertebrate BUSCOs are complete in protein mode (S 95.8 %, D 1.9 %, F 1.4 %, M 1.0 %), against 99.3 % in the genome, so the annotation captures almost all of the conserved gene content the assembly holds. 22,136 genes (89.0 %) have an eggNOG-mapper annotation (Pfam 20,957; GO 15,241; KEGG 15,259); 90.4 % have heart or lung RNA-seq reads and 75.9 % at least ten. Mean gene length is 33.7 kb with 9.8 exons per transcript (mean exon 170 bp, intron 4.4 kb), mean CDS 1,668 bp and median protein 380 aa; 4,355 transcripts are single-exon. The default TSEBRA output would have given 17,392 genes and 89.6 % BUSCO completeness; the comparison of candidate sets is in Supplementary Table S10.

**Figure 3.**
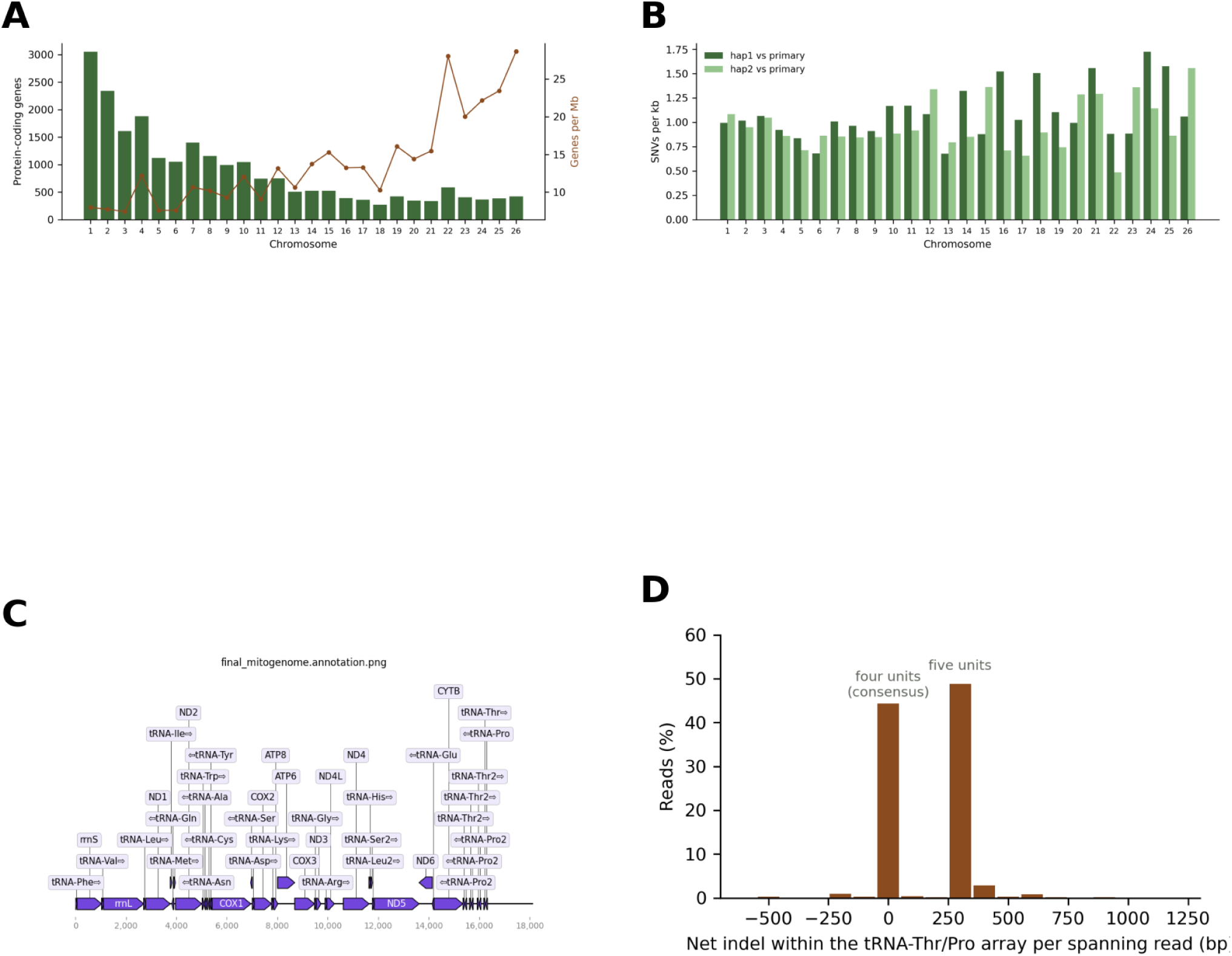
Annotation, haplotypes and mitogenome. (A) Protein-coding genes per chromosome (bars) and gene density (line). (B) SNV density per chromosome between each haplotype assembly and the primary. (C) Mitochondrial genome (18,133 bp, circular; 13 protein-coding genes, 22 tRNAs, 2 rRNAs). (D) Length heteroplasmy of the tRNA-Thr/Pro tandem array: net indel length within the array per spanning HiFi read (0 bp = four-unit consensus, +300 bp = five units).

### Relation to the preliminary version

All 400 sequences of the preliminary version .1 align to Mimp_v3 (≥ 90 % of length for every chromosome-scale sequence); its 32 chromosome-scale sequences correspond to the 26 chromosomes of Mimp_v3 (four of them in two or three pieces), and 56 of its unplaced sequences now lie within chromosomes. Supplementary Table S4 gives the sequence-level correspondence for coordinate transfer.

## Re-use potential

The assembly is a reference for read mapping, variant calling and population and conservation genomics of *M. impressa* and, as the first chromosome-level genome of the basal testudinid genus *Manouria*, an outgroup for comparative studies across Testudinidae. The two haplotype assemblies allow allele-specific analyses and structural-variant discovery within one individual; the correspondence tables map every haplotype scaffold to its primary chromosome. Users of the preliminary version .1 can transfer coordinates with Supplementary Table S4 (chromosome numbers differ between versions; Mimp_v3 numbers chromosomes by size). Mimp_v3 chr12 corresponds to two *G. evgoodei* chromosomes. Regions flagged as heterozygous tandem duplications should be treated with caution in variant calling. The annotation, together with the RNA-seq data from heart and lung, supports expression and comparative studies; the chitin-degradation gene family of this fungivore is one immediate application: the eight-gene GH18 chitinase cluster previously located on version-.1 chr3 lies on Mimp_v3 chr4 (139.05–139.26 Mb, minus strand), where all eight loci now have gene models (MIMP3_g08468–g08479: the four tandem CHIA-clade copies g08472–g08475, the GT31-type gene g08476, the CHI3L-clade and CHIT1-clade genes g08478–g08479, and the formerly fused version-.1 model 088402 resolved into three genes g08468–g08470). The scripted workflow, with its quantitative curation statistics, is reusable for other HiFi + Hi-C genomes.

## Availability of source code and requirements

- Project name: Manouria_impressa_genome_v3 (part of the Chitin_Metabolism_Analysis repository)
- Project home page: https://github.com/snailQH/Manouria_impressa_genome_v3 (public on publication of the Data Note; a Zenodo archive of the release will be linked there)
- perating system: Linux (Slurm cluster); containers via Singularity/Apptainer Programming language: Bash (Slurm scripts), Python 3
- ther requirements: eight conda environments, exported as YAML files (scripts/genome_v3/environments/*.yml, also in Data file 13); containers braker3.sif (BRAKER 3.0.8), tetools.sif (RepeatModeler 2.0.5 / RepeatMasker 4.1.7-p1), fcs-adaptor.sif (0.5.4), fcs- gx.sif (0.5.5), mitohifi.sif (3.2.1)
- License: MIT

## Availability of supporting data and materials

The raw sequencing data are available from the NGDC Genome Sequence Archive (BioProject PRJCA038476): PacBio HiFi reads CRR1746698 and Hi-C reads CRR1746699 (GSA CRA024571; BioSample SAMC4973488, liver), and RNA-seq reads from heart (CRR1747397; BioSample SAMC4973532) and lung (CRR1747398; BioSample SAMC4976828) in GSA CRA024586. The reference assembly Mimp_v3 and its annotation are being released as version 2 of the NGDC Genome Warehouse record GWHHIVU00000000 (version .1, available now, is a preliminary assembly of the same data); the two haplotype assemblies and the mitochondrial genome are being deposited under the same BioProject, and their accessions will be given in the next version of this preprint and in the Data Note.

The complete pipeline, the protocol log, the curation evidence, the QC tables and the scripts that generate every table and figure are in the project repository (https://github.com/snailQH/Manouria_impressa_genome_v3) and will be archived on Zenodo with the Data Note; until the repository is public they are available from the corresponding author.

## Declarations

### List of abbreviations

AGP, A Golden Path; BUSCO, Benchmarking Universal Single-Copy Orthologs; CDS, coding sequence; EBP, Earth BioGenome Project; GSA, Genome Sequence Archive; GWH, Genome Warehouse; HiFi, high-fidelity; Hi-C, high-throughput chromosome conformation capture; MAPQ, mapping quality; NGDC, National Genomics Data Center; PCG, protein-coding gene; QV, consensus quality value; SNV, single-nucleotide variant; SV, structural variant; TE, transposable element.

### Ethics approval and consent to participate

This study was reviewed and approved by the Academic Ethics Committee of the School of Life Sciences, Hainan Normal University, under the Research Ethics and Safety Review Form for Scientific Projects (Approval No. 202532). The animal was collected in Yunnan Province under permit from the Yunnan Provincial Forestry Department (Yunlin Conservation Permit [2013] No. 86, 云林保护许准〔2013〕86号) and the Yunnan Daweishan National Nature Reserve; it died of natural causes in captivity, and all tissues were taken post mortem. No procedure was performed on a living animal.

## Consent for publication

Not applicable.

## Competing interests

The authors declare that they have no competing interests.

## Funding

This work was supported by the National Natural Science Foundation of China (31401983), the Higher Education Scientific Research Project of the Hainan Provincial Department of Education (Hnky2023ZD-5) and the Scientific Research Start-up Fund of Hainan Normal University. The funders had no role in study design, data analysis, interpretation or writing.

## Acknowledgements

We thank the Yunnan Provincial Forestry Department and the Yunnan Daweishan National Nature Reserve for the collection permit, and Annoroad Gene Technology for library preparation and sequencing.

## Supplementary tables

S1 chromosome table (length, contigs, gaps, *G. evgoodei* homolog, preliminary-version name, telomeres); S2 curation edits with Hi-C evidence; S3 Hi-C library statistics; S4 correspondence between the preliminary version .1 and Mimp_v3 (with its assembly statistics); S5 telomere arrays; S6 haplotype → primary correspondence; S7 coverage anomalies; S8 haplotype divergence summary; S9 repeat classes; S10 gene-set statistics and candidate-set comparison; S11 sequences removed by FCS-GX; S12 synteny blocks, chromosome composition and breakpoint evidence relative to *G. evgoodei*. All generated by `scripts/genome_v3/make_manuscript_figs_tables.py` from `results/inputs/`.

---

**Supplementary Figure S1.**
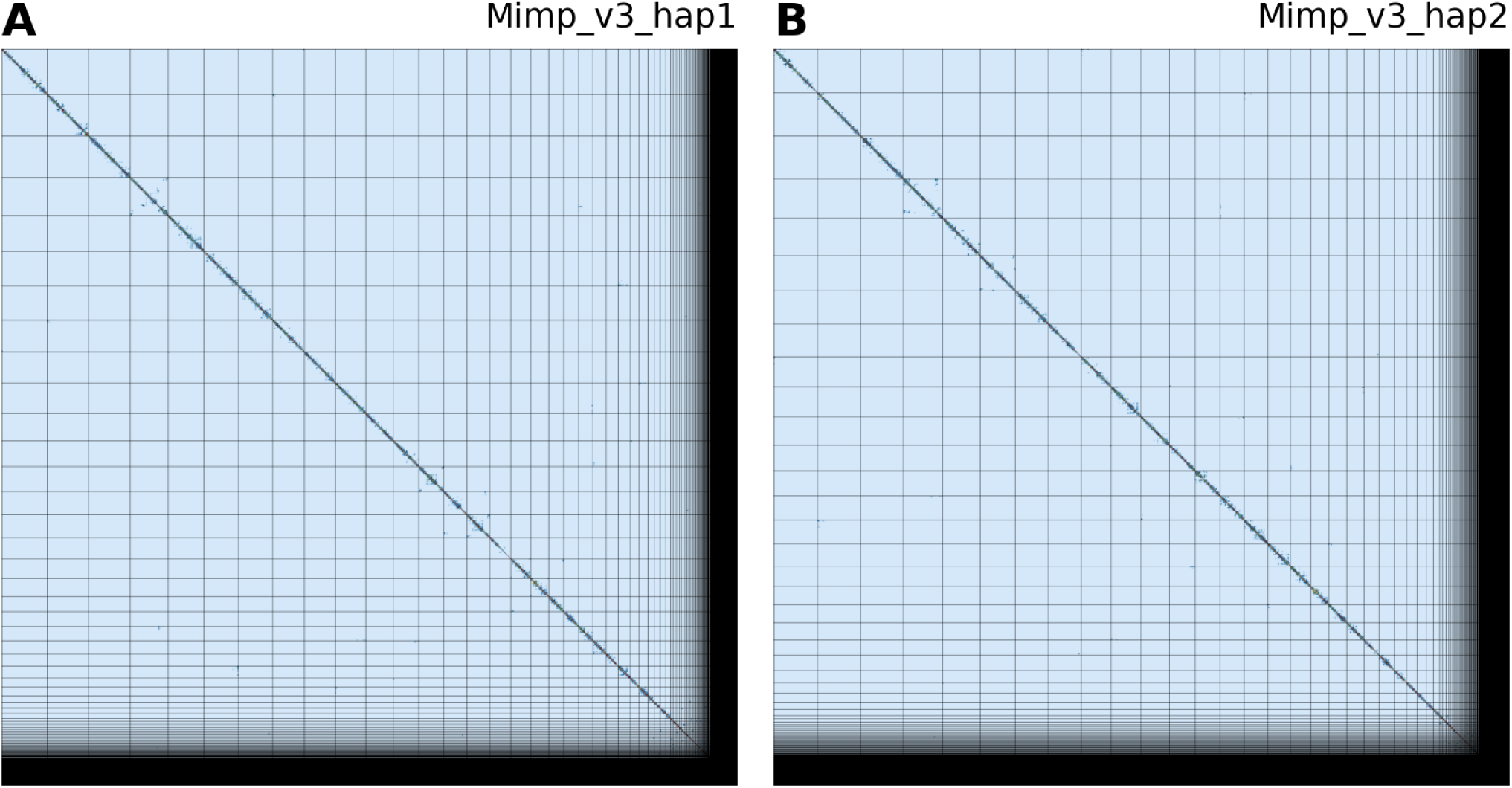
Hi-C contact maps of the two haplotype assemblies (PretextMap).

## Notes

### Competing Interest Statement

The authors have declared no competing interest.

### Summary of Updates

Section on the assembly updated to a newer version. Figures are update based on new assembly, including two haplotypes.

